# Network-specific metabolic cost of functional connectivity in the human brain

**DOI:** 10.64898/2026.09.09.750304

**Authors:** Sarah M Tüchler, Pia Falb, Christian Milz, Matej Murgaš, Murray B Reed, Samantha Graf, Gabriel Schlosser, Clemens Schmidt, Alexandra Mayerweg, Sebastian Klug, Ivan Pörnbacher, Lukas Artmeier, Adam Harouak, Aurelia Sahl, Maximilian Grohmann, Lukas Nics, Godber M Godbersen, Sazan Rasul, Dan Rujescu, Marcus Hacker, Rupert Lanzenberger, Andreas Hahn, the Alzheimer’s Disease Neuroimaging Initiative

## Abstract

Despite decades of extensive research, the complex architecture underlying human brain signaling remains incompletely understood. Previous work investigating the relationship between brain glucose metabolism and functional connectivity (FC) employed whole-brain analysis, without accounting for network interactions, dependencies or hierarchies. Here, we assess network-specific differences in the relationship between FC and metabolic demand, using [^18^F]FDG PET/MR data from three independent datasets. The metabolic cost of FC, i.e., the change in glucose demand associated with corresponding changes in FC, was modeled at rest, during cognitive task performance and in Alzheimer’s disease (AD). Our findings reveal network-specific differences in metabolic cost, with the default mode (DM), somatomotor (SM) and fronto-temporal networks accounting for highest, intermediate and lowest metabolic demands, respectively. Similarly, time-resolved variability of FC demonstrated highest costs for states with DM network involvement and lowest for SM network participation. This relationship was reversed in participants with AD, who exhibited decreased demands in the DM network and increased costs in the SM network. Cognitive performance consistently revealed cost reductions in the DM and SM networks, as well as increases in task relevant networks. Together, these results highlight the flexibility of functional network architecture associated with cognitive demands and neuropathology, and shed light on the complex interplay between glucose metabolism and network interactions.

## INTRODUCTION

Brain function relies on efficient signaling across brain regions, facilitated through dynamically coordinated activity that adapts according to situational requirements (1, 2). These patterns of synchronization are most frequently studied using functional connectivity (FC) (3). FC is derived from functional magnetic resonance imaging (fMRI) by measuring statistical relationships between fMRI blood oxygen level dependent (BOLD) signal time series. Numerous studies have revealed large-scale brain networks of functionally coupled regions (4, 5) and uncovered meaningful information on network organization during rest (6), for various task conditions (7–10) and divergences associated with disease (11, 12). Optimal functioning of this complex system comes with substantial energy demands in terms of glucose consumption (13). Although assessment of the energetic underpinnings of network interactions has produced a wide range of cost-related metrics (14–19), most of these efforts have focused on fMRI alone, often applying a topological or information theoretic notion of energy, instead of metabolic demands. Positron emission tomography (PET) with [^18^F]Fluorodeoxyglucose ([¹⁸F]FDG) provides a biophysiological viewpoint on brain metabolism. Recent advances in functional PET (fPET) imaging (20) further enable the acquisition of time-resolved information on glucose demand during task performance. Previous work investigating the relationship between glucose metabolism and FC at rest (21–24) mostly employed whole-brain analyses without considering network cross-talk. This, however, represents an essential aspect, as brain network topology is organized through network interactions, dependencies and hierarchies (25–27). Similar to spatial differentiation, this has also been established for the temporal domain. It is widely recognized that the so-called resting-state actually comprises different transient states (28–30), characterized by distinct, recurring FC patterns, indicative of varying signaling architecture (29), which are often referred to as dynamic FC states. Finally, changes in network connectivity have also been reported across a variety of task conditions (7, 8, 31–34), in aging (35, 36) and in various brain disorders, such as in Alzheimer’s disease (AD) (37) or Schizophrenia (11, 38). However, the relationship between these FC changes and glucose metabolism has not been sufficiently investigated, especially with regard to network interactions.

To address this knowledge gap, we introduce an integrative approach that explicitly assesses glucose expenditure associated with FC, using simultaneously acquired BOLD fMRI and [¹⁸F]FDG PET data. Our multimodal approach allows joint estimation of network-specific metabolic cost, as well as the relative contribution of distinct networks to metabolic demands. This analysis is further extended to FC variability over time, by determining dynamic states and associated differences in metabolic cost. This is realized in two independent datasets of healthy young adults performing different cognitive tasks, as well as healthy elderly subjects and patients with AD from the ADNI cohort.

## MATERIALS & METHODS

### Study design

For the following analyses data from two different studies were used, both of which included resting-state and task conditions, as described below (datasets 1 and 2). For a more detailed description please refer to the respective manuscripts (33, 34). Participants underwent one or multiple simultaneous PET/MR scans at the Department of Radiology and Nuclear Medicine at the Medical University of Vienna with a hybrid PET/MR system (Biograph mMR, Siemens Healthineers, Erlangen, Germany), equipped with a 12-channel head coil. Additional data stem from the ADNI 2 dataset, which includes PET and fMRI measurements from healthy elderly participants and subjects with AD. These were obtained from the Alzheimer’s Disease Neuroimaging Initiative (ADNI) database (adni.loni.usc.edu). The ADNI was launched in 2003 as a public-private partnership, led by Principal Investigator Michael W. Weiner, MD. The primary goal of ADNI has been to test whether serial magnetic resonance imaging (MRI), positron emission tomography (PET), other biological markers, and clinical and neuropsychological assessment can be combined to measure the progression of mild cognitive impairment (MCI) and early Alzheimer’s disease (AD). The current goals include validating biomarkers for clinical trials, improving the generalizability of ADNI data by increasing diversity in the participant cohort, and to provide data concerning the diagnosis and progression of Alzheimer’s disease to the scientific community. For up-to-date information, see adni.loni.usc.edu.

For dataset 1 (DS1), subjects performed a modified version of the videogame Tetris, as outlined previously (34). The task was repeated at different levels of difficulty (easy and hard), however, for the current analysis only resting-state and task data of the ‘hard’ condition were used. Each measurement started with the acquisition of structural MRI sequences, followed by a 52-minute fPET measurement for which the radiotracer [^18^F]FDG was applied using a bolus (1 minute) plus constant infusion (51 minutes) protocol (5.1 Mbq/kg body weight, 20% as bolus). Tracer administration was carried out with a perfusion pump (Syramed µSP6000, Arcomed, Regensdorf, Switzerland) and synthesis of the radiotracer [^18^F]FDG was performed as described previously (20). The measurement consisted of an 8-minute resting-state baseline followed by four 6-minute task blocks, which were interleaved with 5-minute periods of rest. During task performance BOLD data were acquired simultaneously.

For dataset 2 (DS2) participants underwent an adapted version of the Montral Imaging Stress Test (MIST) (39), during which subjects performed mathematical calculations in two different versions: first without any stress-inducing elements and then again with a set time limit and negative feedback on their performance, designed to induce psychosocial stress (33, 39). To ensure individual stress levels of the participants were not unduly influenced by outside factors, such as anxiety due to the unknown environment, each measurement started with a one-hour acclimatization period in the gantry that included MRI and low-level cognitive inputs.

This was followed by 56 minutes of fPET with the radioligand [^18^F]FDG, applying a bolus (1 minute) plus constant infusion (55 minutes) protocol (5.1 MBq/kg body weight, 20% as bolus). fPET and BOLD fMRI were acquired simultaneously during control (no feedback or time limit) and stress blocks (feedback and time limit). Each task block lasted for 8 minutes and the two conditions were separated by 9 minutes of rest. To enable absolute quantification of the cerebral metabolic rate of glucose (CMRGlu), a cardiac image-derived input function was acquired between task blocks and supplemented by venous blood samples (40).

The ADNI 2 study protocol included separate scanning sessions of structural and BOLD fMRI imaging, as well as [^18^F]FDG PET data acquisition at resting-state, for which [^18^F]FDG was applied as a single bolus.

### Participants

The datasets comprise 51 healthy participants for DS1 (mean age ± SD = 23.3 ± 3.3 years, 24 female) and 56 healthy participants for DS2 (mean age ± SD = 23.9 ± 4.9, 28 female). All subjects underwent an initial screening visit, during which a standard medical examination was performed, including blood tests, electrocardiography, neurological testing and the Structural Clinical Interview for DSM-IV (DS1) or DSM-V (DS2). Additionally, a urine pregnancy test was administered at the screening visit and before the PET/MR examination to rule out pregnancy. Exclusion criteria comprised pregnancy or breastfeeding, contraindications to MRI, as well as limits for previous research-related radiation exposure. Participants experiencing somatic, neurological or psychiatric disorders, currently or within the last 12 months, and with current or past substance abuse or psychotropic medication were also excluded. For DS1, subjects with previous Tetris experience within the last 3 years were excluded. Only right-handed subjects between 18 and 40 years were considered for these studies.

From the ADNI 2 cohort data from 42 healthy elderly participants (mean age ± SD = 76.3 ± 7.1, 22 female) and 26 subjects who were assigned to the Early-Stage-Alzheimer’s-Disease cohort (mean age ± SD = 72.8 ± 7.8, 12 female) were used.

### MRI data acquisition

Measurements for DS1 and DS2 were carried out on a Siemens Biograph mMR scanner.

The structural MRI for DS1 and DS2 was acquired with a T1-weighted MPRAGE sequence (TE/TR = 4.21/2200 ms, matrix size = 240 × 256, 160 slices, voxel size = 1 × 1 × 1 mm + 0.1 mm gap, 7.7 min). For both DS1 and DS2 BOLD data were acquired with an EPI sequence (TE/TR = 30/2000 ms, matrix size = 80 × 80, 34 slices, voxel size = 2.5 × 2.5 x 2.5 mm + 0.825 mm gap).

From the ADNI dataset a T1-weighted MPRAGE sequence (TE/TR = 3.16/6800 ms, flip angle = 9°, voxel size = 1.2 × 1 × 1 mm) and BOLD fMRI at resting state (TE/TR = 30/3000 ms, flip angle = 80°, voxel size = 3.3125 × 3.3125 × 3.313 mm, 7 min) are available.

### Blood sampling

As discussed in previous work (20), DS1 implemented manual arterial blood sampling 3, 4, 5, 14, 25, 36 and 47 minutes after the initial tracer application to determine the arterial input function.

DS2 applied a different approach (40, 41), relying on a cardiac image-derived input function and supplementary venous samples, which were collected at minutes 19.5, 36 and 50.5 after the initial tracer application.

The whole-blood- and plasma activity were measured with a gamma counter (Wizard2, 3”, PerkinElmer, Waltham, MA, USA) and subsequent multiplication of the whole-blood arterial or image-derived input function with the average plasma-to-whole-blood ratio yielded the final input function. The ADNI dataset did not include any blood sampling.

### Quantification of CMRGlu

fPET data of DS1 and DS2 were reconstructed to 30 s frames (matrix size = 344×344, 127 slices) with an Ordinary Poisson Ordered Subset Expectation Maximization Algorithm (OP-OSEM), and attenuation correction was performed using a database approach (42, 43). Image preprocessing was performed in SPM12 (https://www.fil.ion.ucl.ac.uk/spm/) and included motion correction, spatial normalization via the T1-weighted structural MRI, and smoothing with a Gaussian kernel of 8 mm. Analysis of both datasets was carried out with the fPET toolbox (44) using a general linear model with 4 regressors, which were used to distinguish between task effects and baseline uptake. DS1 comprised one baseline (mean time course of average gray matter voxels, excluding regions with significant BOLD changes during the hard task condition (34)) and two task regressors (easy and hard, linear ramp functions), as well as an additional regressor for head motion. DS2 also used a baseline regressor (average across gray matter voxels, excluding voxels involved in stress processing as identified in a meta-analysis (45)), two task regressors (control and stress, linear ramp functions) and head motion.

The influx constants Ki for baseline and tasks were determined with the Gjedde-Patlak plot, and then used to calculate the cerebral metabolic rate of glucose (CMRGlu) with a lumped constant of 0.89 (46) and glucose plasma levels as obtained before the scan (average of three values).

For the ADNI dataset data was acquired in 6 frames and standardized uptake value ratio (SUVR) with reference to whole-brain gray matter was used for further analysis.

### Computation of static and dynamic functional connectivity

Preprocessing of fMRI data was carried out in SPM12 as described previously (20) and included slice time and motion correction, after which data were spatially normalized via the T1-weighted image and smoothed with an 8 mm Gaussian kernel. Motion scrubbing was performed using the standardized per-image standard deviation of the temporal derivative (zDVARS) (47). Regression analysis was carried out to remove potentially confounding signals originating from white matter, cerebrospinal fluid and motion with subsequent bandpass filtering (0.01 < f < 0.15 Hz) (34, 48).

For this analysis 216 regions were assessed in cortical (200 parcels) and subcortical (16 parcels) areas, as previously defined by Schaefer et al (49) and Tian et al (50), respectively. Functional connectivity was determined at rest for all datasets and during task performance for DS1 (8 minutes resting-state and 6 minutes task) and DS2 (8 minutes resting-state, control and stress condition).

First, static FC was calculated using Pearson’s correlation coefficient between each region’s BOLD time course and the resulting matrices were transformed using Fisher’s r-to-z-transformation.

To evaluate dynamic FC changes in resting-state over time, an exponentially tapered sliding window approach was used to segment the full BOLD time series of each region of interest into overlapping windows (51). Each dynamic window had a duration of 60 s (51) and was weighted using an exponential function (52):

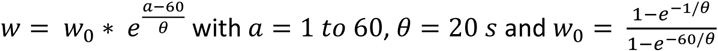

The window was advanced by 1 frame (2 s), at each iteration. These parameters were chosen in accordance with relevant literature (52–54). Finally, Fisher’s r-to-z-transformation was applied to the resulting dynamic FC matrices.

### Identification and characterization of dynamic functional connectivity states

To identify dynamic states, windows from dynamic FC calculations were thresholded at 20% (51). Subsequent clustering with the kmeans clustering algorithm (Correlation distance, 10 replicates, a maximum of 200 iterations, Matlab 2018 and 2023) was performed to identify distinct and recurring patterns of FC and the cluster count was assessed by applying the elbow criterion to the within/between sum of differences. The resulting clusters could then be interpreted as functional brain states that reflect temporal variations of connectivity arising from the heterogeneity of resting-state connectivity (30).

In order to characterize these dynamic states, their total occurrence over all subjects and windows was calculated, as was the prevalence of transitions between them.

Furthermore, connectograms visualizing FC across and within networks were created with Circos (55). To improve interpretability and reduce influence of artifactual correlations, each state’s 216x216 FC matrix was thresholded at 10% and all remaining elements were assigned to the corresponding resting-state brain networks (DA, dorsal attention; DM, default mode; FP, frontoparietal; FT, frontotemporal; SM, somatomotor; VA, ventral attention; VI, visual) (4) and the basal ganglia (B)).

### Metabolic cost of static functional connectivity at rest

Association between static FC and CMRGlu demand across brain networks was assessed with linear regression analysis separately for each subject (Figure 1). Network specific regressors were created that represent a region’s connectivity strength with each network. This was done by averaging each row of the 216x216 FC matrix over regions of a particular network, yielding a matrix of 216 regions x 8 networks as independent variables. The dependent variable was the average CMRGlu per region (vector of 216 regions). The beta estimates of this model reflect the unit change of CMRGlu per unit change of FC within each network. We use the term metabolic cost as an operational descriptor of this association, rather than as a direct measure of energetic expenditure caused by connectivity. As FC strength of all networks is included in a single model, this reflects network-specific differences in metabolic cost of FC while accounting for collinearity between networks. Of note, the metabolic demand per FC has to be interpreted relative to the overall brain metabolism, thus a negative value implies that the cost is less than average.

**Figure 1.**
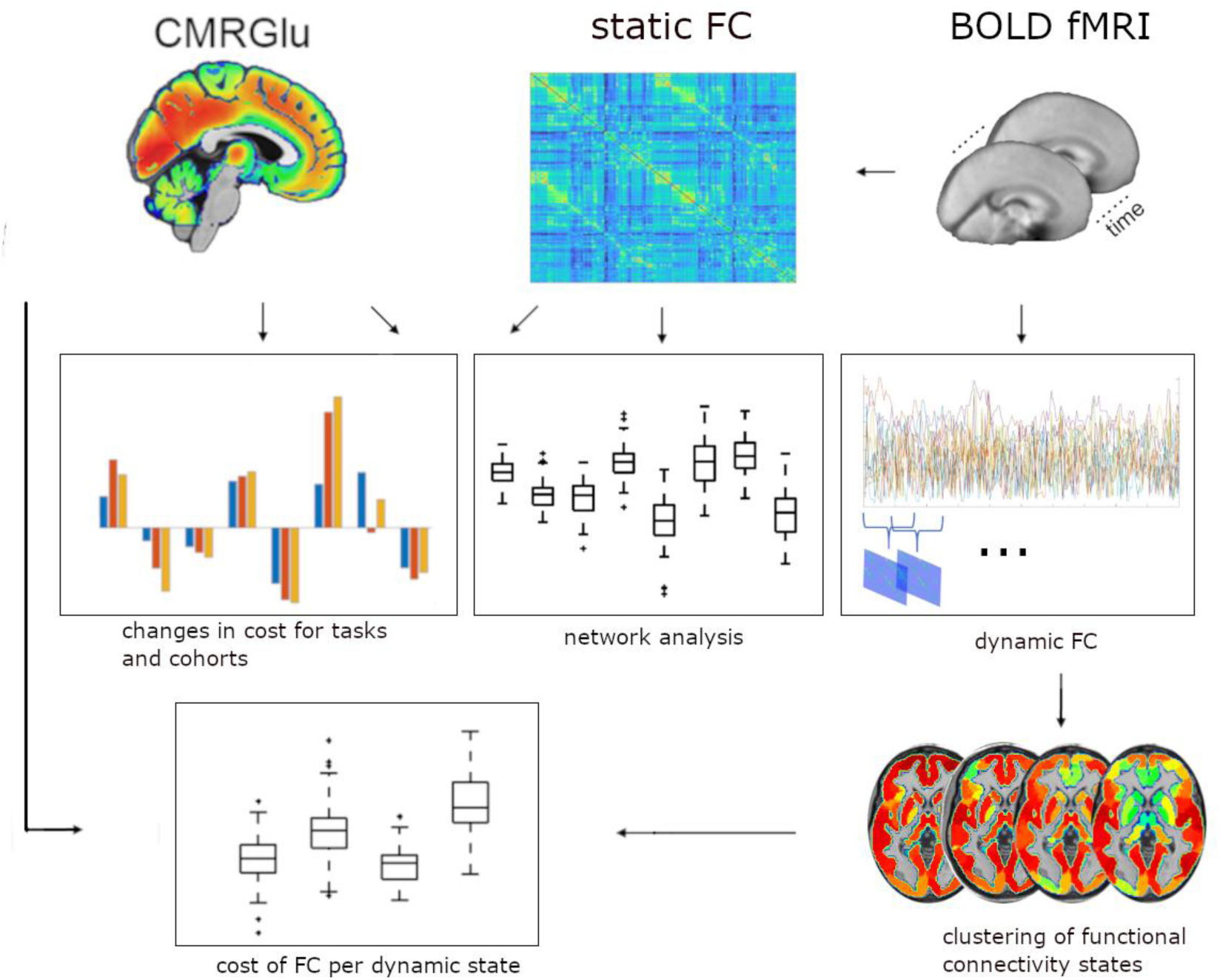
Overview of analytical process: Blood oxygen level dependent (BOLD) functional magnetic resonance imaging (fMRI) data is used to calculate static and dynamic functional connectivity (FC). Differences in the relationship between static FC and cerebral metabolic rate of glucose (CMRGlu) across networks are analyzed. Dynamic FC windows are clustered into distinct dynamic states, enabling calculation of the metabolic cost for FC states and transitions between them. Additionally, changes in metabolic cost of FC during task performance and in subjects with AD are investigated using static FC and CMRGlu/SUVR. CMRGlu and BOLD fMRI illustrations in the upper row of this figure were taken from (56) under CC BY license.

### Metabolic cost of functional connectivity in dynamic states

Association of dynamic states and transitions with CMRGlu demand across brain networks was assessed with linear regression analysis separately for each subject, analogous to the above procedure. First, the dynamic FC maps of each cluster and subject were averaged, resulting in four FC states (see results), i.e., four 216x216 matrices that depict the average state-specific FC pattern of a subject. Then, mean connectivity strength of each region in a state was determined through averaging each row of the four FC matrices across subjects, producing a state-based regressor that captures a region’s connectivity strength for each state (independent variables of 216 regions x 4 states). Again, the dependent variable was the average CMRGlu per region (vector of 216 regions). The computed beta estimates of this model therefore reflect the unit change of CMRGlu per unit change of FC within each state and capture state-specific differences in metabolic cost of FC.

Similarly, the metabolic cost of transitioning between different states was evaluated with a general linear model. Connectivity differences for each subject were obtained by subtracting the average FC matrix of the origin state from the target state’s average FC matrix. This resulted in 6 transitions and averaging of rows yielded a matrix of 216 regions x 6 transitions as the independent variable.

### Effects of task performance, aging and AD

To investigate task-related changes in metabolic cost of static FC, the above analysis was repeated for subjects of DS1 and DS2 during task performance. The average CMRGlu of each region was extracted and then modeled by static FC during task performance using a general linear model with a matrix of 216 regions x 8 networks as independent variable, separately for each task. Finally, effects of AD on the metabolic cost of static FC were assessed. Since CMRGlu maps were not available for the ADNI dataset, SUVR was used instead, keeping the above analysis pipeline otherwise identical. Even though CMRGlu and SUVR are related quantities, as they both reflect glucose metabolism, their respective regression coefficients cannot be readily compared. Therefore, SUVR maps were also extracted for DS1 and DS2, and resting-state analysis was repeated using SUVR to evaluate potential differences.

### Statistical analysis

As mentioned above, the resulting beta estimates from the regression models represent the unit change of CMRGlu (or SUVR) per unit change of FC within each network, state or transition, for each subject. To assess if each network’s metabolic cost of resting-state FC differs from the overall average, one sample t-tests were used. Paired t-tests were performed to test if the metabolic cost of FC differs between rest and task conditions. Similarly, differences between healthy elderly subjects and AD patients were assessed using two-sample t-tests. Bonferroni Holm correction was used to account for multiple comparisons, respectively (8 networks, 4 states, 6 transitions).

## RESULTS

### Metabolic cost of functional connectivity during rest

The metabolic cost of FC was determined as the change of CMRGlu per unit change of FC. Our findings show significant network-specific differences, with the DM and FT networks accounting for highest and lowest metabolic demands, respectively (Figure 2 a-b), in both DS1 and DS2. Still, apart from the SM network in DS1, metabolic costs of FC reached significance for all functional networks (p< 0.05 corrected for multiple comparisons) and qualitative comparison showed high agreement between both datasets.

**Figure 2.**
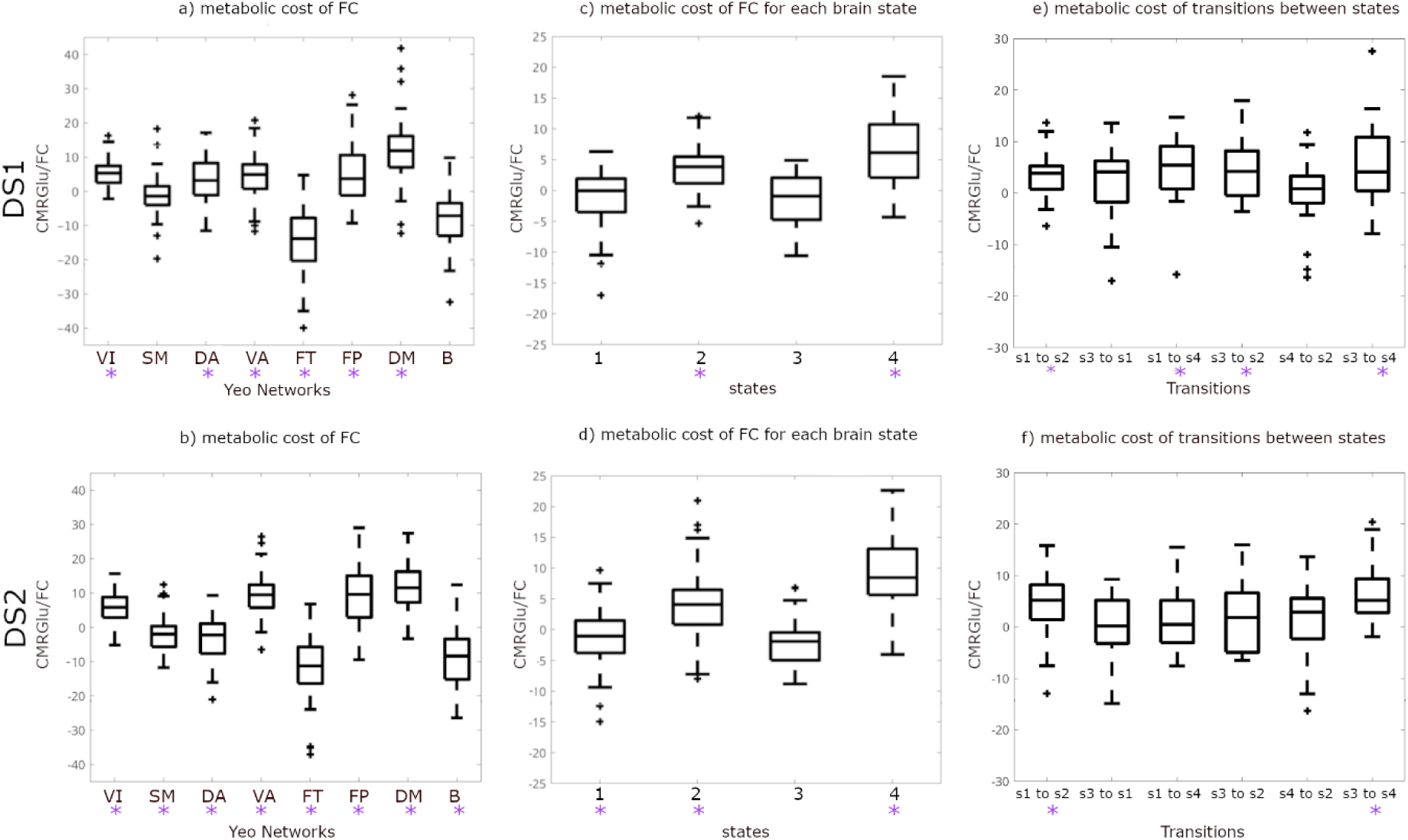
Overview of metabolic cost of functional connectivity (FC) for networks, states and transitions of DS1 and DS2: a - b) Metabolic cost of static resting-state FC across different networks. c - d) Metabolic cost of FC for each brain state, which were defined through kmeans clustering of dynamic FC windows. e - f) Metabolic cost associated with transitions between different dynamic states. *p<0.05 corrected for multiple comparisons. DA, dorsal attention; DM, default mode; FP, frontoparietal; FT, frontotemporal; SM, somatomotor; VA, ventral attention; VI, visual; B, basal ganglia.

### Metabolic Cost of functional connectivity in different brain states

The kmeans clustering approach revealed four distinct states, which is in the range of previously reported state counts (28, 57–59). These four states were consistently identified over DS1 and DS2, and states of both datasets were matched by maximizing correlation between their respective centroids (supp fig 1). Glucose demand per unit change of FC differed substantially across dynamic states, with two states showing significantly high metabolic costs in both DS1 and DS2 (p<0.05 corrected for multiple comparisons, Figure 2 c-d).

Since connectivity patterns fluctuate between these dynamic states, we also analyzed the expense associated with state transitions. Transitions from state 1 to 2 and from state 3 to 4 were particularly metabolically expensive for both datasets, with DS1 also exhibiting significant cost for transition 1 to 4 and 3 to 2 (p<0.05 corrected for multiple comparisons, Figure 2e-f). Notably, transitions from states with lower metabolic demand per FC to those with high costs were consistently associated with higher transition expenditure (Figure 2c-f).

### Analysis of connectivity states

Given these variations between states, we aimed to further assess their specific FC, number of occurrences and rate of transitions between them. Evaluation of state-specific static FC showed good agreement between DS1 and DS2 (Figure 3a) and revealed inherent differences between states. State 1 exhibited fairly high FC within the DM and SM networks, as well as a strong link between the SM and VA networks. State 2 was characterized by DA connections, with high within-network FC and links to FP, SM and VA networks. State 3 showed high connectivity of the SM network with the DA and VA networks, and high within-network FC. State 4 exhibited distinctive characteristics with strong links between the DM and FP networks, along with strong within-network connectivity of the DM network. Unlike the other states, state 4 displayed only a minor link between SM and VA networks, as well as low within-network FC of the SM network. The high cost in state 4 is in line with our findings from static FC, which also identified the DM network as metabolically most expensive.

**Figure 3.**
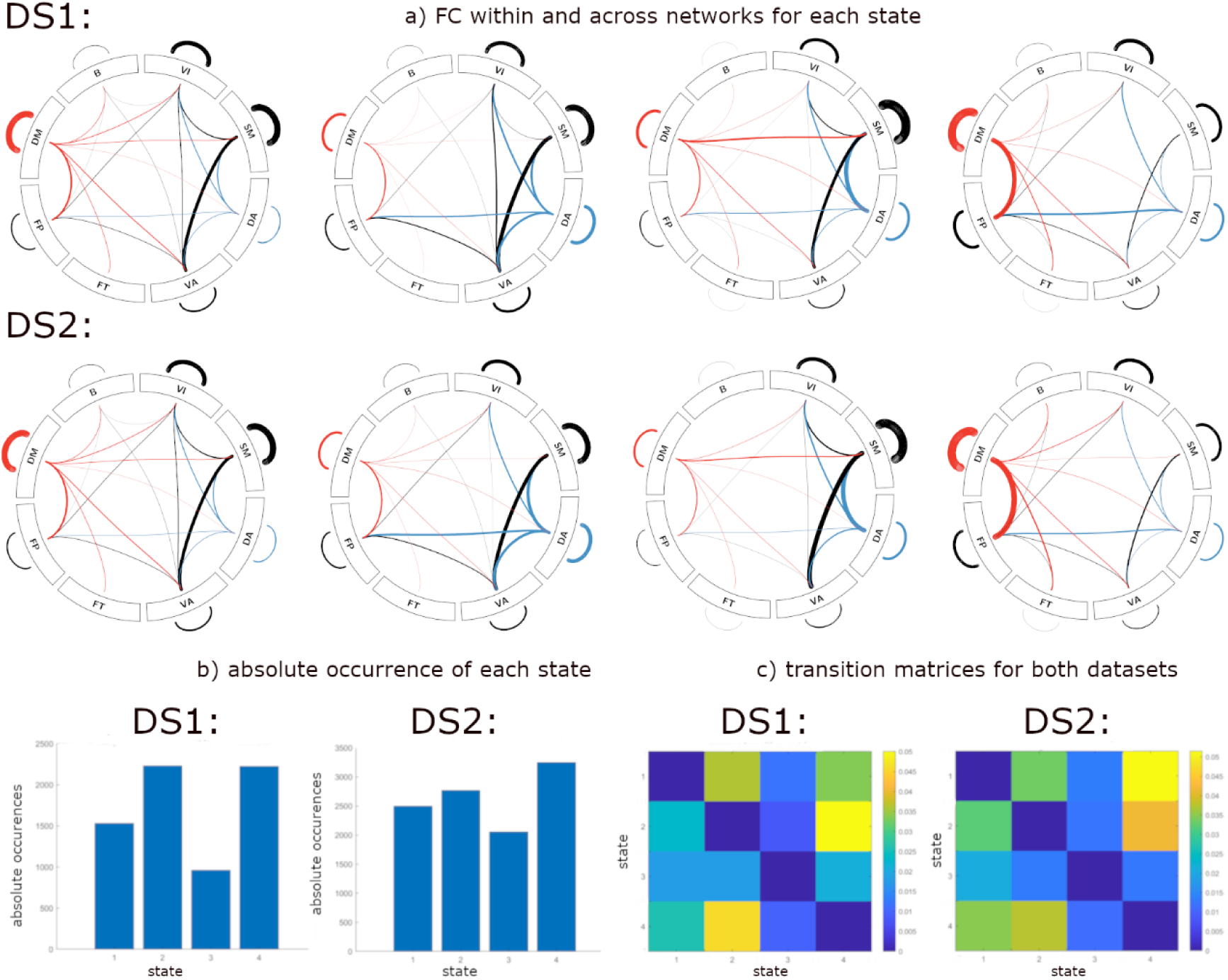
Characterization of brain states: a) Circos (55) representation of functional connectivity (FC) within and across networks for all four states of each dataset. Links with the DM network are colored red and links with the DA network are in blue for better visibility. Link thickness represents strength of the connection. FC matrices were thresholded to a 10% level. b) Absolute occurrence counts of each state, summed over all subjects of a dataset. c) Transition matrices, where each row represents the probability of a state transitioning into another. Diagonal was set to 0 for better interpretability due to high probability to remain in a state. DA, dorsal attention; DM, default mode; FP, frontoparietal; FT, frontotemporal; SM, somatomotor; VA, ventral attention; VI, visual; B, basal ganglia.

Interestingly, those states with highest metabolic demands (states 2 and 4) also showed highest occurrence rates (Figure 3c). It should be noted that analysis of state timelines revealed subjects which did not occupy all four states over the course of their measurement, which was especially prevalent for state 3. Consistent with high occurrence rates, the transition matrices (Figure 3b) showed high probabilities for transitions into state 2 and state 4.

### Effects of task performance and aging

Changes in metabolic cost of FC in different task conditions as compared to resting-state (Figure 4 a-c) showed good agreement particularly for SM and DM networks over DS1 and DS2, despite differing task paradigms. However, also task-specific changes were observed, especially in DA and FP networks, which are involved in processing of the respective tasks. Specifically, for DS1, differences between task performance and resting-state were observed particularly in DA, SM and DM networks (figure 4a). Subjects of DS2 showed significant differences in metabolic cost for both task conditions in SM, FP and DM networks, and additionally in the VI network for the control condition (figure 4b). However, no significant differences for the comparison control vs stress emerged.

**Figure 4.**
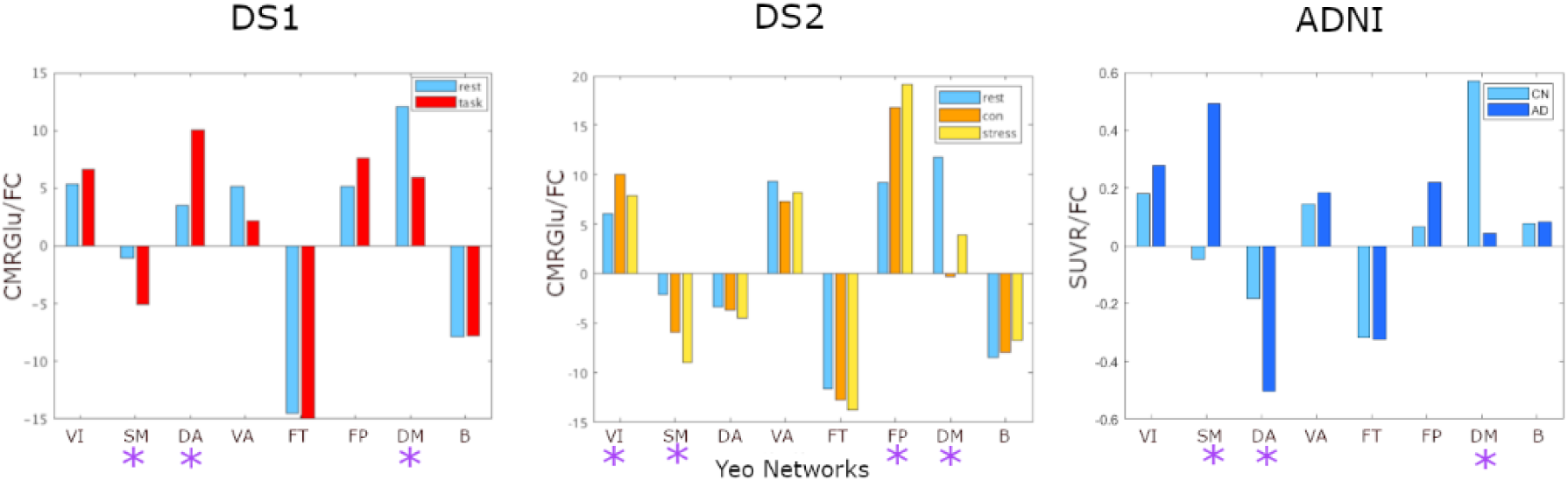
Changes in metabolic cost of FC during rest (blue), task performance (yellow, orange and red), with age (light blue) and in AD (dark blue) across networks: a) Differences between metabolic cost of FC during performance of a Tetris-based task vs rest emerged in SM, DA and DM networks. b) Task-specific differences in metabolic cost of FC for MIST performance during control and stress conditions were observed in VI, SM, FP and DM networks (control vs rest), and in SM, FP and DM networks (stress vs rest), no significant differences occurred comparing control and stress. c) Differences in metabolic cost of FC between patients with AD and a control group of healthy elderly subjects; significant differences emerged in SM, DA and DM networks. * p<0.05 corrected for multiple comparisons. DA, dorsal attention; DM, default mode; FP, frontoparietal; FT, frontotemporal; SM, somatomotor; VA, ventral attention; VI, visual; B, basal ganglia.

Healthy elderly subjects of the ADNI dataset showed similar results to our young cohorts apart from subcortical regions. Comparisons between the ADNI dataset (SUVR) and the results of DS1 and DS2 (CMRGlu) were enabled by repeating the latter analyses with SUVR, which showed great similarity across both metrics (Supp Fig 1). Comparing AD patients to healthy elderly controls revealed a significant decrease in metabolic cost in the DM and DA networks, along with significant increases in the SM network (figure 4c).

## DISCUSSION

Using simultaneous PET/MR imaging we estimated the metabolic cost of FC across cortical and subcortical networks and investigated changes related to task performance, healthy aging and AD. At rest, DM and FT networks emerged with highest and lowest metabolic demands, respectively. During cognitive performance, these costs were redistributed with general changes in SM and DM as well as task-specific changes in DA and FP networks as recruited by the respective cognitive load. We further identified four distinct dynamic FC states, two of which exhibited above-average metabolic costs and were characterized by strong involvement of the DA or DM networks. Investigation of AD-related changes in metabolic costs of FC further revealed decreases in the DA and DM networks, highlighting the particular vulnerability of these networks in this patient cohort.

### Metabolic cost of resting-state FC

Our analysis of intrinsic resting-state FC identified the DM and FP networks as regions with high metabolic cost of FC. The DM network is thought to be active during rest and is implicated in a large number of self-referential processes, such as mind-wandering and introspective processing (60), whereas the FP network is hypothesized to be vital for flexible reconfiguration of task states according to external demands (61, 62). The pattern we observed might reflect the increased flexibility necessary to adapt between internally-directed cognition, attention modulation and preparedness to capture external stimuli (14) or reconfigure network structures to enable goal-directed behavior (61, 63). This is in line with previous findings which located regions with higher energetic cost of signaling predominantly in the FP and DM networks (21). Interestingly, these high metabolic demands of the DM and FP control networks are also mirrored in regional differences of aerobic glycolysis, which is expressed at high levels in regions of the DM network and in regions linked to task control processes, contrasting with low levels in sensory cortices (64). Moreover, networks with higher- or lower-than-average metabolic cost were mostly transmodal networks, whereas unimodal sensory networks showed less pronounced differences (26, 65). Together, this may indicate that higher-order networks use fast-acting energy production via aerobic glycolysis (66) to facilitate rapid switches between internal and external cognitive demands.

These static FC findings were further manifested in the analysis of dynamic connectivity states, where states with pronounced involvement of higher order networks also demonstrated increased metabolic demands. State 4 in particular, the most metabolically expensive state, was characterized by high FC within the DM network and between DM and FP networks, indicating high synchronization. This combined involvement of DM and FP networks is in line with established signaling architectures involved in autobiographical planning (61, 67). Conversely, state 3 with strong involvement of SM network showed low metabolic cost. Surprisingly, the distinct metabolic demands associated with state 4 (high cost) and 3 (low cost) were further accompanied by the highest and lowest occurrence rates, respectively, and by corresponding costs associated with state transitions from cheap to expensive states in the in both analyzed datasets. This possibly indicates a potential trade-off between cost and efficiency, mirroring fundamental constraints in structural connectivity (15, 68, 69). It is worth highlighting that similar effects were observed for two completely independent datasets. This applies to static FC as well as dynamic FC states, with strong agreement of states, transitions and the resulting metabolic costs. Considering the distinct organization of dynamic states (32, 70) along with links to general intelligence (71), psychological disorders (58), and cognitive decline (36), investigation of their metabolic demands may reveal further insights regarding cost-efficiency trade-offs of hierarchical network organization.

### Task-induced changes in metabolic cost

Subsequent analysis of the different cognitive tasks and their associated metabolic cost of FC also revealed regional variations. Overall, comparison between DS1 and DS2 resulted in high similarity of task-induced changes in metabolic demands of FC, despite differing cognitive processes, particularly in the DM and SM networks (decreases), FP network (increases) as well as FT network and B (stable). However, also task-specific differences were observed.

The initial analysis of the Tetris-based task (DS1) showed task-evoked increases of CMRGlu predominantly in VI and DA networks, along with consistent negative responses in the DM network (8, 34). These changes were largely reflected by metabolic costs of FC, with significant increases and decreases in the DA and DM networks respectively. Performance of the cognitively challenging Tetris task requires active visuo-spatial reasoning (34), which corresponds to VI and DA network activations, potentially increasing costs to grant higher neuronal flexibility necessary for dynamic task-induced adaptations.

A similar, though task-specific, pattern was observed for the stress task (DS2). Here, CMRGlu increases in the FP network during the control condition (33) were accompanied by increased metabolic cost of FC. However, a notable divergence stood out for the DM network. Although FC’s metabolic demand decreased (similar to DS1), CMRGlu remained the same for the control condition or even increased during psychosocial stress (33) (in contrast to DS1). This may be related to correspondingly altered FC during stress, which decreased within the DM and increased within the FP network (33). In sum, our current assessment highlights the distinct ratio of changes in CMRGlu and FC. Interestingly, while the comparison between rest and control/stress conditions produced substantial differences, the metabolic costs of FC did not change significantly between control and stress condition, despite the reported functional and metabolic stress response. This indicates that during stress, overall FC and glucose uptake increase concordantly, keeping the relative cost constant.

### Age related effects

Considering the known effect of AD on network organization (37, 72) and metabolism (73), we investigated potential pathological alterations of associations between these parameters.

The DM network has been shown to be particularly vulnerable to amyloid-beta deposition (72, 74), which in turn is associated with hypometabolism and cognitive decline (75). Variations of FC patterns with age emerge from complex dynamic reorganizations of functional brain networks (76, 77). For example, a number of previous studies have reliably identified the posterior DM network as a region of decreased FC in AD, while recent work has found increased connectivity in the frontal DM network regions in early stages of the disease (78–80). Additionally, evidence has suggested decreased connectivity across multiple networks in healthy aging (81). AD causes further decreases in within-network connectivity of DM and DA networks, indicative of network degradation (82, 83), along with decreased structural integration and segregation (84, 85), ultimately contributing to an overall decline of functionality. Glucose metabolism as captured by [^18^F]FDG PET follows a similar pattern, showing characteristic decreases in neocortical association areas including the posterior cingulate, precuneus, temporoparietal and frontal multimodal association regions, whereas the sensorimotor and visual cortices are largely spared (86).

Our findings showed distinct changes in metabolic cost of FC in the DM, DA and SM networks between healthy elderly subjects and patients with AD. Since both CMRGlu and FC in the DM network are known to decrease with disease progression, the reported cost reduction suggests that glucose demand is more strongly reduced than FC. Conversely, increased demands in the SM network point toward a reverse relationship. Interestingly, metabolic costs in healthy elderly compared to young subjects revealed cost increases in the subcortex, providing insights into the desynchrony of metabolic and functional decline.

In sum, the DM network exhibits high metabolic demands (87) and harbors a particularly high number of hubs (72), which are characterized by their extensive connectivity and are thought to be essential for functional integration (88). Thus, the association between FC and CMRGlu in the DM network may offer an approach to assess AD-related effects and potential alterations in the coupling between FC and glucose metabolism.

### Limitations, conclusions and future work

Our work proposes a strategy to investigate the underlying metabolic demands of brain FC. However, several limitations need to be acknowledged. Previous work (22) has provided reports of non-linear effects in the relationship between glucose metabolism and FC, suggesting plateauing per-degree-costs for high connectivity. Since we apply a general linear model, any potential non-linear interactions are not taken into consideration. However, previous work did not consider network interactions, but rather assessed networks independently, or brain connectivity as a whole. Furthermore, it should be mentioned that FC only reflects synchronicity of BOLD signal time courses across regions and does not account for other factors such as wiring efficiency, which might be a confounder of metabolic cost and affect our results (23). Thus, future work could attempt to integrate structural connectivity information into the proposed model of metabolic cost.

In conclusion, our approach demonstrated that overall metabolic cost of both static and dynamic FC was stable across two independent datasets and showed network specific differences in response to certain cognitive tasks and in an AD cohort. As FC and glucose metabolism are interdependent but distinct entities, this may offer a promising avenue for further evaluation in disease cohorts.

## ACKNOWLEDGMENTS

We thank the graduated team members and the diploma students of the Neuroimaging Lab (NIL, head R. Lanzenberger) as well as the clinical colleagues from the Department of Psychiatry and Psychotherapy for clinical and/or administrative support.

Data collection and sharing for the Alzheimer’s Disease Neuroimaging Initiative (ADNI) is funded by the National Institute on Aging (National Institutes of Health Grant U19 AG024904). The grantee organization is the Northern California Institute for Research and Education. In the past, ADNI has also received funding from the National Institute of Biomedical Imaging and Bioengineering, the Canadian Institutes of Health Research, and private sector contributions through the Foundation for the National Institutes of Health (FNIH) including generous contributions from the following: AbbVie, Alzheimer’s Association; Alzheimer’s Drug Discovery Foundation; Araclon Biotech; BioClinica, Inc.; Biogen; Bristol-Myers Squibb Company; CereSpir, Inc.; Cogstate; Eisai Inc.; Elan Pharmaceuticals, Inc.; Eli Lilly and Company; EuroImmun; F. Hoffmann-La Roche Ltd and its affiliated company Genentech, Inc.; Fujirebio; GE Healthcare; IXICO Ltd.; Janssen Alzheimer Immunotherapy Research & Development, LLC.; Johnson & Johnson Pharmaceutical Research &Development LLC.; Lumosity; Lundbeck; Merck & Co., Inc.; Meso Scale Diagnostics, LLC.; NeuroRx Research; Neurotrack Technologies; Novartis Pharmaceuticals Corporation; Pfizer Inc.; Piramal Imaging; Servier; Takeda Pharmaceutical Company; and Transition Therapeutics.

## AUTHOR CONTRIBUTIONS (CRediT)

**Sarah M Tüchler**: Investigation, Data curation, Methodology, Formal Analysis, Visualization, Writing – original draft, **Pia Falb**: Investigation, Data curation **Christian Milz**: Investigation, Data curation, Funding acquisition **Matej Murgaš**: Investigation, Data curation **Murray B Reed**: Software, Validation, Data curation **Samantha Graf**: Investigation **Gabriel Schlosser**: Investigation **Clemens Schmidt**: Investigation **Alexandra Mayerweg**: Investigation **Ivan Pörnbacher**: Investigation **Lukas Artmeier**: Investigation **Adam Harouak**: Investigation **Aurelia Sahl**: Project administration **Maximilian Grohmann**: Investigation **Lukas Nics**: Supervision, Project administration **Godber M Godbersen:** Supervision **Sazan Rasul**: Supervision **Dan Rujescu**: Supervision, Resources **Marcus Hacker**: Supervision, Resources **Rupert Lanzenberger**: Conceptualization, Supervision, Resources, Project administration, Funding acquisition **Andreas Hahn**: Conceptualization, Methodology, Software, Validation, Formal analysis, Visualization, Supervision, Funding acquisition, Writing – original draft. **All authors**: Writing - Review & Editing

## DATA AND CODE AVAILABILITY

Raw data will not be publicly available due to reasons of data protection. Processed data and custom code can be obtained from the corresponding author with a data sharing agreement, approved by the departments of legal affairs and data clearing of the Medical University of Vienna.

## ETHICS

DS1 and DS2: After detailed explanation of the study protocol, all participants gave written informed consent. The study was approved by the Ethics Committee (DS1 ethics number: 1479/2015; DS2 ethics number 1642/2022) of the Medical University of Vienna and procedures were carried out in accordance with the Declaration of Helsinki.

DS1: Clinical trial registration ClinicalTrials.gov ID: NCT03485066.

DS2: Clinical trial registration ClinicalTrials.gov ID: NCT06243783

## REFERENCES

1. Deng S, Li J, Thomas Yeo BT, Gu S. Control theory illustrates the energy efficiency in the dynamic reconfiguration of functional connectivity. Commun Biol. 2022;5(1):295.

2. Cocchi L, Zalesky A, Fornito A, Mattingley JB. Dynamic cooperation and competition between brain systems during cognitive control. Trends Cogn Sci. 2013;17(10):493–501.

3. Biswal B, Yetkin FZ, Haughton VM, Hyde JS. Functional connectivity in the motor cortex of resting human brain using echo-planar MRI. Magn Reson Med. 1995;34(4):537–41.

4. Yeo BTT, Krienen FM, Sepulcre J, Sabuncu MR, Lashkari D, Hollinshead M, et al. The organization of the human cerebral cortex estimated by intrinsic functional connectivity. J Neurophysiol. 2011;106(3):1125–65.

5. Power JD, Cohen AL, Nelson SM, Wig GS, Barnes KA, Church JA, et al. Functional network organization of the human brain. Neuron. 2011;72(4):665–78.

6. Rogers BP, Morgan VL, Newton AT, Gore JC. Assessing functional connectivity in the human brain by fMRI. Magn Reson Imaging. 2007;25(10):1347–57.

7. Arbabshirani MR, Havlicek M, Kiehl KA, Pearlson GD, Calhoun VD. Functional network connectivity during rest and task conditions: a comparative study. Hum Brain Mapp. 2013;34(11):2959–71.

8. Godbersen GM, Klug S, Wadsak W, Pichler V, Raitanen J, Rieckmann A, et al. Task-evoked metabolic demands of the posteromedial default mode network are shaped by dorsal attention and frontoparietal control networks. Elife. 2023;12.

9. Shi L, Sun J, Xia Y, Ren Z, Chen Q, Wei D, et al. Large-scale brain network connectivity underlying creativity in resting-state and task fMRI: Cooperation between default network and frontal-parietal network. Biol Psychol. 2018;135:102–11.

10. Yang YL, Deng HX, Xing GY, Xia XL, Li HF. Brain functional network connectivity based on a visual task: visual information processing-related brain regions are significantly activated in the task state. Neural Regen Res. 2015;10(2):298–307.

11. Xu R, Zhang X, Zhou S, Guo L, Mo F, Ma H, et al. Brain structural damage networks at different stages of schizophrenia. Psychol Med. 2024;54(16):4809–19.

12. He Q, Yang Z, Xue B, Song X, Zhang C, Yin C, et al. Epilepsy alters brain networks in patients with insular glioma. CNS Neurosci Ther. 2024;30(6):e14805.

13. Yu Y, Herman P, Rothman DL, Agarwal D, Hyder F. Evaluating the gray and white matter energy budgets of human brain function. J Cereb Blood Flow Metab. 2018;38(8):1339–53.

14. Saberi M, Rieck JR, Golafshan S, Grady CL, Misic B, Dunkley BT, et al. The brain selectively allocates energy to functional brain networks under cognitive control. Sci Rep. 2024;14(1):32032.

15. Avena-Koenigsberger A, Yan X, Kolchinsky A, van den Heuvel MP, Hagmann P, Sporns O. A spectrum of routing strategies for brain networks. PLoS Comput Biol. 2019;15(3):e1006833.

16. Bassett DS, Bullmore ET, Meyer-Lindenberg A, Apud JA, Weinberger DR, Coppola R. Cognitive fitness of cost-efficient brain functional networks. Proc Natl Acad Sci U S A. 2009;106(28):11747–52.

17. Achard S, Bullmore E. Efficiency and cost of economical brain functional networks. PLoS Comput Biol. 2007;3(2):e17.

18. Gu S, Cieslak M, Baird B, Muldoon SF, Grafton ST, Pasqualetti F, et al. The Energy Landscape of Neurophysiological Activity Implicit in Brain Network Structure. Sci Rep. 2018;8(1):2507.

19. Sengupta B, Stemmler MB, Friston KJ. Information and efficiency in the nervous system--a synthesis. PLoS Comput Biol. 2013;9(7):e1003157.

20. Rischka L, Gryglewski G, Pfaff S, Vanicek T, Hienert M, Klobl M, et al. Reduced task durations in functional PET imaging with [(18)F]FDG approaching that of functional MRI. Neuroimage. 2018;181:323–30.

21. Castrillon G, Epp S, Bose A, Fraticelli L, Hechler A, Belenya R, et al. An energy costly architecture of neuromodulators for human brain evolution and cognition. Sci Adv. 2023;9(50):eadi7632.

22. Tomasi D, Wang GJ, Volkow ND. Energetic cost of brain functional connectivity. Proc Natl Acad Sci U S A. 2013;110(33):13642–7.

23. Aiello M, Salvatore E, Cachia A, Pappatà S, Cavaliere C, Prinster A, et al. Relationship between simultaneously acquired resting-state regional cerebral glucose metabolism and functional MRI: A PET/MR hybrid scanner study. Neuroimage. 2015;113:111–21.

24. Palombit A, Silvestri E, Volpi T, Aiello M, Cecchin D, Bertoldo A, et al. Variability of regional glucose metabolism and the topology of functional networks in the human brain. Neuroimage. 2022;257:119280.

25. Fox MD, Snyder AZ, Vincent JL, Corbetta M, Van Essen DC, Raichle ME. The human brain is intrinsically organized into dynamic, anticorrelated functional networks. Proc Natl Acad Sci U S A. 2005;102(27):9673–8.

26. Smallwood J, Bernhardt BC, Leech R, Bzdok D, Jefferies E, Margulies DS. The default mode network in cognition: a topographical perspective. Nat Rev Neurosci. 2021;22(8):503–13.

27. Uddin LQ, Supekar KS, Ryali S, Menon V. Dynamic reconfiguration of structural and functional connectivity across core neurocognitive brain networks with development. J Neurosci. 2011;31(50):18578–89.

28. Allen EA, Damaraju E, Plis SM, Erhardt EB, Eichele T, Calhoun VD. Tracking whole-brain connectivity dynamics in the resting state. Cereb Cortex. 2014;24(3):663–76.

29. Calhoun VD, Miller R, Pearlson G, Adali T. The chronnectome: time-varying connectivity networks as the next frontier in fMRI data discovery. Neuron. 2014;84(2):262–74.

30. Hutchison RM, Womelsdorf T, Allen EA, Bandettini PA, Calhoun VD, Corbetta M, et al. Dynamic functional connectivity: promise, issues, and interpretations. Neuroimage. 2013;80:360–78.

31. Hahn A, Gryglewski G, Nics L, Rischka L, Ganger S, Sigurdardottir H, et al. Task-relevant brain networks identified with simultaneous PET/MR imaging of metabolism and connectivity. Brain Struct Funct. 2018;223(3):1369–78.

32. Vidaurre D, Abeysuriya R, Becker R, Quinn AJ, Alfaro-Almagro F, Smith SM, et al. Discovering dynamic brain networks from big data in rest and task. Neuroimage. 2018;180(Pt B):646–56.

33. Schlosser G, Milz C, Falb P, Graf S, Tüchler SM, Murgaš M, et al. Differential upregulation of metabolic demands and functional integration of the default mode network during stress. BioRxiv 2026.08.14.744771 [Preprint]. 2026 [cited 01.09.2026]. doi: 10.64898/2026.08.14.744771

34. Hahn A, Breakspear M, Rischka L, Wadsak W, Godbersen GM, Pichler V, et al. Reconfiguration of functional brain networks and metabolic cost converge during task performance. Elife. 2020;9.

35. Wu ZX, Gao YY, Potter T, Benoit J, Shen J, Schulz PE, et al. Interactions Between Aging and Alzheimer’s Disease on Structural Brain Networks. Front Aging Neurosci. 2021;13.

36. Cabral J, Vidaurre D, Marques P, Magalhaes R, Silva Moreira P, Miguel Soares J, et al. Cognitive performance in healthy older adults relates to spontaneous switching between states of functional connectivity during rest. Sci Rep. 2017;7(1):5135.

37. Hahn K, Myers N, Prigarin S, Rodenacker K, Kurz A, Forstl H, et al. Selectively and progressively disrupted structural connectivity of functional brain networks in Alzheimer’s disease - revealed by a novel framework to analyze edge distributions of networks detecting disruptions with strong statistical evidence. Neuroimage. 2013;81:96–109.

38. Wang L, Metzak PD, Honer WG, Woodward TS. Impaired efficiency of functional networks underlying episodic memory-for-context in schizophrenia. J Neurosci. 2010;30(39):13171–9.

39. Dedovic K, Renwick R, Mahani NK, Engert V, Lupien SJ, Pruessner JC. The Montreal Imaging Stress Task: using functional imaging to investigate the effects of perceiving and processing psychosocial stress in the human brain. J Psychiatry Neurosci. 2005;30(5):319–25.

40. Reed MB, Handschuh PA, Schmidt C, Murgas M, Gomola D, Milz C, et al. Validation of cardiac image-derived input functions for functional PET quantification. Eur J Nucl Med Mol I. 2024;51(9):2625–37.

41. Reed MB, Godbersen GM, Vraka C, Rausch I, Ponce de Leon M, Popper V, et al. Comparison of cardiac image-derived input functions for quantitative whole body [(18)F]FDG imaging with arterial blood sampling. Front Physiol. 2023;14:1074052.

42. Burgos N, Cardoso MJ, Thielemans K, Modat M, Pedemonte S, Dickson J, et al. Attenuation correction synthesis for hybrid PET-MR scanners: application to brain studies. IEEE Trans Med Imaging. 2014;33(12):2332–41.

43. Milz CM, M. Merida, I. Silberbauer, L.R. Nics, L. Godbersen, G.M. Gryglewski, G. Hacker, M. Costes, N., Hammers A. Open-access template and database approaches for pseudo-CT generation in brain PET/MRI attenuation correction. 2025.

44. Hahn A, Reed MB, Milz C, Falb P, Murgas M, Lanzenberger R. A unified approach for identifying PET-based neuronal activation and molecular connectivity with the functional PET toolbox. J Cereb Blood Flow Metab. 2026;46(2):558–68.

45. Qiu Y, Fan Z, Zhong M, Yang J, Wu K, Huiqing H, et al. Brain activation elicited by acute stress: An ALE meta-analysis. Neurosci Biobehav Rev. 2022;132:706–24.

46. Hahn A, Gryglewski G, Nics L, Hienert M, Rischka L, Vraka C, et al. Quantification of Task-Specific Glucose Metabolism with Constant Infusion of 18F-FDG. J Nucl Med. 2016;57(12):1933–40.

47. Afyouni S, Nichols TE. Insight and inference for DVARS. Neuroimage. 2018;172:291–312.

48. Sun FT, Miller LM, D’Esposito M. Measuring interregional functional connectivity using coherence and partial coherence analyses of fMRI data. Neuroimage. 2004;21(2):647–58.

49. Schaefer A, Kong R, Gordon EM, Laumann TO, Zuo XN, Holmes AJ, et al. Local-Global Parcellation of the Human Cerebral Cortex from Intrinsic Functional Connectivity MRI. Cereb Cortex. 2018;28(9):3095–114.

50. Tian Y, Margulies DS, Breakspear M, Zalesky A. Topographic organization of the human subcortex unveiled with functional connectivity gradients. Nat Neurosci. 2020;23(11):1421–32.

51. Zalesky A, Fornito A, Cocchi L, Gollo LL, Breakspear M. Time-resolved resting-state brain networks. Proc Natl Acad Sci U S A. 2014;111(28):10341–6.

52. Pozzi F, Di Matteo T, Aste T. Exponential smoothing weighted correlations. Eur Phys J B. 2012;85(6).

53. Hahn A, Strandberg TO, Stomrud E, Nilsson M, van Westen D, Palmqvist S, et al. Association Between Earliest Amyloid Uptake and Functional Connectivity in Cognitively Unimpaired Elderly. Cereb Cortex. 2019;29(5):2173–82.

54. Shirer WR, Ryali S, Rykhlevskaia E, Menon V, Greicius MD. Decoding subject-driven cognitive states with whole-brain connectivity patterns. Cereb Cortex. 2012;22(1):158–65.

55. Krzywinski M, Schein J, Birol I, Connors J, Gascoyne R, Horsman D, et al. Circos: an information aesthetic for comparative genomics. Genome Res. 2009;19(9):1639–45.

56. Klug S, Murgas M, Godbersen GM, Hacker M, Lanzenberger R, Hahn A, et al. Synaptic signaling modeled by functional connectivity predicts metabolic demands of the human brain. Neuroimage. 2024;295:120658.

57. Goldhacker M, Tome AM, Greenlee MW, Lang EW. Frequency-Resolved Dynamic Functional Connectivity Reveals Scale-Stable Features of Connectivity-States. Front Hum Neurosci. 2018;12:253.

58. Damaraju E, Allen EA, Belger A, Ford JM, McEwen S, Mathalon DH, et al. Dynamic functional connectivity analysis reveals transient states of dysconnectivity in schizophrenia. Neuroimage Clin. 2014;5:298–308.

59. Patanaik A, Tandi J, Ong JL, Wang C, Zhou J, Chee MWL. Dynamic functional connectivity and its behavioral correlates beyond vigilance. Neuroimage. 2018;177:1–10.

60. Menon V. 20 years of the default mode network: A review and synthesis. Neuron. 2023;111(16):2469–87.

61. Spreng RN, Stevens WD, Chamberlain JP, Gilmore AW, Schacter DL. Default network activity, coupled with the frontoparietal control network, supports goal-directed cognition. Neuroimage. 2010;53(1):303–17.

62. Marek S, Dosenbach NUF. The frontoparietal network: function, electrophysiology, and importance of individual precision mapping. Dialogues Clin Neurosci. 2018;20(2):133–40.

63. Zuo NM, Yang ZY, Liu Y, Li J, Jiang TZ. Both activated and less-activated regions identified by functional MRI reconfigure to support task executions. Brain Behav. 2018;8(1).

64. Vaishnavi SN, Vlassenko AG, Rundle MM, Snyder AZ, Mintun MA, Raichle ME. Regional aerobic glycolysis in the human brain. Proc Natl Acad Sci U S A. 2010;107(41):17757–62.

65. Margulies DS, Ghosh SS, Goulas A, Falkiewicz M, Huntenburg JM, Langs G, et al. Situating the default-mode network along a principal gradient of macroscale cortical organization. P Natl Acad Sci USA. 2016;113(44):12574–9.

66. Theriault JE, Shaffer C, Dienel GA, Sander CY, Hooker JM, Dickerson BC, et al. A functional account of stimulation-based aerobic glycolysis and its role in interpreting BOLD signal intensity increases in neuroimaging experiments. Neurosci Biobehav Rev. 2023;153:105373.

67. Spreng RN, Schacter DL. Default network modulation and large-scale network interactivity in healthy young and old adults. Cereb Cortex. 2012;22(11):2610–21.

68. Ma J, Zhang J, Lin Y, Dai Z. Cost-efficiency trade-offs of the human brain network revealed by a multiobjective evolutionary algorithm. Neuroimage. 2021;236:118040.

69. Bullmore E, Sporns O. The economy of brain network organization. Nat Rev Neurosci. 2012;13(5):336–49.

70. Betzel RF, Fukushima M, He Y, Zuo XN, Sporns O. Dynamic fluctuations coincide with periods of high and low modularity in resting-state functional brain networks. Neuroimage. 2016;127:287–97.

71. Ng J, Yu JC, Feusner JD, Hawco C. Higher general intelligence is associated with stable, efficient, and typical dynamic functional brain connectivity patterns. Imaging Neurosci (Camb). 2024;2.

72. Buckner RL, Sepulcre J, Talukdar T, Krienen FM, Liu H, Hedden T, et al. Cortical hubs revealed by intrinsic functional connectivity: mapping, assessment of stability, and relation to Alzheimer’s disease. J Neurosci. 2009;29(6):1860–73.

73. Mosconi L. Brain glucose metabolism in the early and specific diagnosis of Alzheimer’s disease. FDG-PET studies in MCI and AD. Eur J Nucl Med Mol Imaging. 2005;32(4):486–510.

74. Mormino EC, Smiljic A, Hayenga AO, Onami SH, Greicius MD, Rabinovici GD, et al. Relationships between beta-amyloid and functional connectivity in different components of the default mode network in aging. Cereb Cortex. 2011;21(10):2399–407.

75. Pascoal TA, Mathotaarachchi S, Kang MS, Mohaddes S, Shin M, Park AY, et al. Abeta-induced vulnerability propagates via the brain’s default mode network. Nat Commun. 2019;10(1):2353.

76. Malagurski B, Deschwanden PF, Jancke L, Merillat S. Longitudinal functional connectivity patterns of the default mode network in healthy older adults. Neuroimage. 2022;259:119414.

77. Malagurski B, Liem F, Oschwald J, Merillat S, Jancke L. Longitudinal functional brain network reconfiguration in healthy aging. Hum Brain Mapp. 2020;41(17):4829–45.

78. Vemuri P, Jones DT, Jack CR, Jr. Resting state functional MRI in Alzheimer’s Disease. Alzheimers Res Ther. 2012;4(1):2.

79. Damoiseaux JS, Prater KE, Miller BL, Greicius MD. Functional connectivity tracks clinical deterioration in Alzheimer’s disease. Neurobiol Aging. 2012;33(4):828 e19-30.

80. Jones DT, Machulda MM, Vemuri P, McDade EM, Zeng G, Senjem ML, et al. Age-related changes in the default mode network are more advanced in Alzheimer disease. Neurology. 2011;77(16):1524–31.

81. Deery HA, Di Paolo R, Moran C, Egan GF, Jamadar SD. The older adult brain is less modular, more integrated, and less efficient at rest: A systematic review of large-scale resting-state functional brain networks in aging. Psychophysiology. 2023;60(1):e14159.

82. Chhatwal JP, Schultz AP, Johnson KA, Hedden T, Jaimes S, Benzinger TLS, et al. Preferential degradation of cognitive networks differentiates Alzheimer’s disease from ageing. Brain. 2018;141(5):1486–500.

83. Brier MR, Thomas JB, Snyder AZ, Benzinger TL, Zhang D, Raichle ME, et al. Loss of intranetwork and internetwork resting state functional connections with Alzheimer’s disease progression. J Neurosci. 2012;32(26):8890–9.

84. Liu W, Zuo C, Chen L, Lan H, Luo C, Li X, et al. The whole-brain structural and functional connectome in Alzheimer’s disease spectrum: A multimodal Bayesian meta-analysis of graph theoretical characteristics. Neurosci Biobehav Rev. 2025;174:106174.

85. Cassady KE, Adams JN, Chen X, Maass A, Harrison TM, Landau S, et al. Alzheimer’s Pathology Is Associated with Dedifferentiation of Intrinsic Functional Memory Networks in Aging. Cereb Cortex. 2021;31(10):4781–93.

86. Herholz K, Carter SF, Jones M. Positron emission tomography imaging in dementia. Br J Radiol. 2007;80 Spec No 2:S160-7.

87. Shah NJ, Arrubla J, Rajkumar R, Farrher E, Mauler J, Kops ER, et al. Multimodal Fingerprints of Resting State Networks as assessed by Simultaneous Trimodal MR-PET-EEG Imaging. Sci Rep-Uk. 2017;7.

88. Sporns O, Zwi JD. The small world of the cerebral cortex. Neuroinformatics. 2004;2(2):145–62.

