## Supplementary figures 1 and 2 for "Network-specific metabolic cost of functional connectivity in the human brain"

### SUPPLEMENTARY MATERIALS

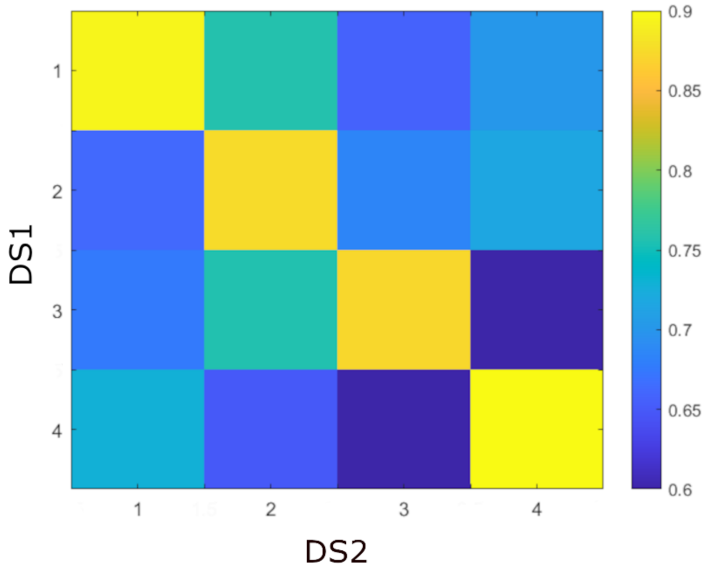


*Supplementary Figure 1: Correlation matrix of states' centroids between datasets*

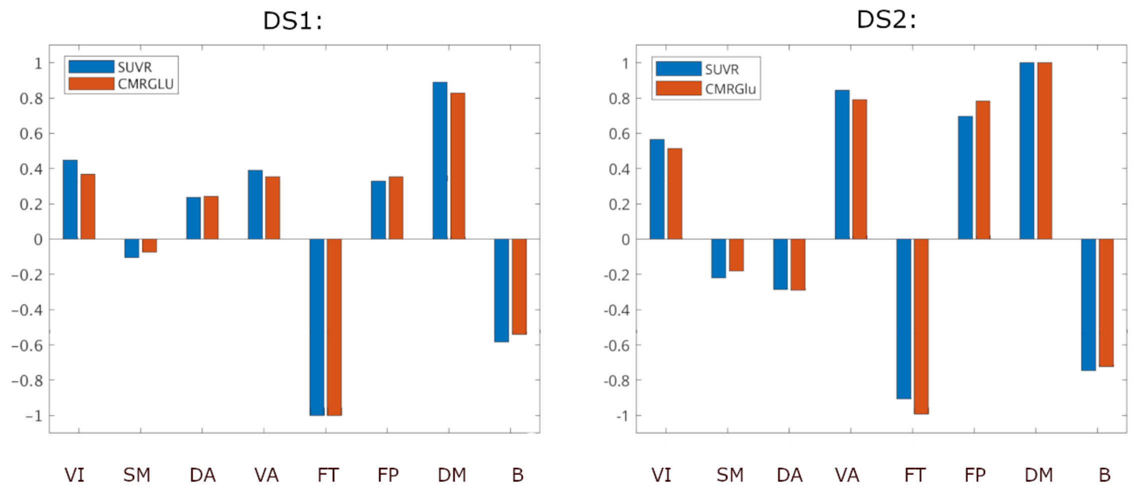


*Supplementary Figure 2: Normalized resting-state metabolic costs of FC calculated in terms of CMRGLU and SUVR to enable comparison*
